# Measuring the net outcome of conditional mutualism: a case study with oaks and scatterhoarding rodents

**DOI:** 10.1101/232264

**Authors:** Michał Bogdziewicz, Elizabeth E. Crone, Rafał Zwolak

**Affiliations:** Department of Systematic Zoology, Faculty of Biology, Adam Mickiewicz University, Poznań, Poland; CREAF, Cerdanyola del Valles, Catalonia, 08193 Spain; Department of Biology, Tufts University, 163 Packard Ave, Medford, MA 02155, USA; Department of Environmental Science and Policy, University of California at Davis, One Shields Avenue, Davis, CA 95616

**Keywords:** antagonism, biotic invasion, conditional mutualism, scatterhoarding, seed dispersal

## Abstract

Numerous interactions between plants and animals vary in their outcome between antagonism and mutualism, but it has proven to be difficult to quantify their final outcome. Interactions between plants and scatterhoarding animals provide a prime example of this phenomenon. Scatterhoarders consume large quantities of seeds (potentially reducing plant establishment), yet also disperse seeds and bury them in shallow caches (potentially improving recruitment). However, it has been rarely determined which role prevails for particular plant species. We demonstrate how plant-scatterhoarder interactions can be placed at the antagonism-mutualism continuum, with interactions between rodents and two oaks species (sessile oak *Quercus petraea*, and red oak *Q. rubra)* as an empirical example. Our approach consists of quantifying the net outcome of the interaction through assembling different vital rates (e.g. probability of seedling recruitment with and without rodents; near and far from conspecific trees; with and without seed pilferage) piecewise with a simple mathematical model. Our results indicate that during the period of the study, interactions between scatterhoarding rodents and both focal oaks were antagonistic. Even though acorn burial increased the likelihood of seedling establishment, this effect was not strong enough to compensate for the costs of seed predation. Furthermore, we found no evidence that the short-distance transportation that is usually provided by small mammals benefited early oak recruitment. Our study demonstrates how readily accessible field data can be used to gauge the outcomes in conditional mutualisms.

## Introduction

Numerous interactions between plants and animals vary in their outcome between antagonism and mutualism (Bronstein, 1994; Palmer *et al*., 2010). Interactions between plants and scatterhoarding animals, such as rodents or corvids, are a prime example of this phenomenon because scatterhoarders play a dual role in plant regeneration. On the one hand, they consume large quantities of seeds and reduce plant establishment (Howe & Brown, 2001; Zwolak *et al*., 2010; Larios *et al*., 2017). On the other hand, however, they disperse seeds and bury them in shallow caches, which for some plant species provide the only means of successful recruitment (Vander Wall, 1992; Asquith *et al*., 1999; Muñoz & Bonal, 2011; Pesendorfer *et al*., 2016). Determining which role prevails for particular plant species can help guide nature conservation, forest management, and control of invasive species because management strategies will depend on the interaction outcome. Yet, these outcomes have been rarely quantified.

Whether scatterhoarding granivores are beneficial or detrimental for plant population depends on whether recruitment with granivores is greater or less than recruitment without seed caching (Jansen & Forget, 2001; Theimer, 2005; Schupp *et al*., 2010; Zwolak & Crone, 2012). Quantifying this has proven challenging, but one approach is to build up the net outcome from separately-measured components (Zwolak & Crone, 2012). This kind of approach is similar to predicting population dynamics from separately-measured vital rates (e.g., Morris & Doak 2002), but has been used less often to evaluate species interactions. Intuitively, when the benefits from seed caching are high, plants can bear higher costs in the form of seed consumption. Following on this notion, the interaction is mutualistic when the probability of caching and not retrieving cached seeds exceeds the ratio of seedling emergence from surface to seedling emergence from caches.

To briefly review our past use of this approach (Zwolak & Crone, 2012), we started from the premise that granivores are beneficial when plant recruitment in the presence of granivores is greater than plant recruitment in the absence of granivores. This inequality is written in mathematical terms as follows:

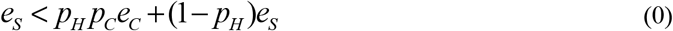

Where *e_S_* is seedling emergence from surface, *p_H_* is the proportion of seeds harvested by granivores, *p_C_* is the probability that seeds will be cached and left uneaten, and *e_C_* is the seedling emergence from caches. In studies of plant-granivore interactions, it is much easier and more common to measure the emergence rates (*e_S_* and *e_C_*) than the caching rates (p_H_ and *p_C_*; see Zwolak and Crone 2012). Therefore, in order to compare the role of granivores across studies, Zwolak and Crone (2012) rearranged the equation to calculate the minimum value of *p_C_* that would be necessary for granivores to increase plant recruitment:

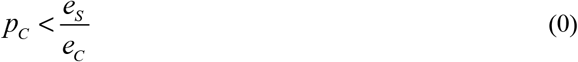

Thus, granivores help plant recruitment when the proportion of cached and uneaten seeds exceeds a threshold value (hereafter *p_C_*, after Zwolak and Crone 2012) determined by the seedling emergence ratio, i.e. the benefits of seed burial.

Scatterhoarders not only bury the seeds in the topsoil, but also move them away from the parent plant. This also modifies recruitment probability, with effects that are usually believed to be positive, due to colonization of ephemeral microsites or escape from distance- and density dependent mortality (Jansen *et al*., 2008; Comita *et al*., 2010; Johnson *et al*., 2012; Fricke *et al*., 2014). However, the effects can also be negative, e.g. when habitat quality is autocorrelated, it often declines with distance from maternal plants (John *et al*., 2007; Condit *et al*., 2013). Furthermore, the distance to the seed source may alter rodent foraging activity and seed pilferage rates through changes in local seed availability (Stapanian & Smith, 1984; Gálvez *et al*., 2009). Nonetheless, even though factors shaping dispersal distance by scatterhoarders, especially by rodents, are intensively studied (Jansen *et al*., 2004; Xiao *et al*., 2005; Moore *et al*., 2007; Sunyer *et al*., 2014; Lichti *et al*., 2017), the actual influence of dispersal distance on recruitment probability is seldom quantified. The intertwined escape-related (transportation distance) and condition-related (burial) benefits constrain our ability to understand mechanisms that drive the ecological interactions between plants and scatterhoarders.

Here, we use empirical data to illustrate an approach for separating burial- and distance-dependent benefits of rodent seed dispersal. We used two oak species as model system: sessile oak *(Quercus petraea)*, and northern red oak (Q. *rubra*). The sessile oak is the dominant native oak in Central European forests. The northern red oak was introduced to European forests from North America in the 17th century as an ornamental species (Woziwoda *et al*., 2014b). Currently, it is one of the most frequent foreign deciduous species in the region (Woziwoda *et al*., 2014b), and its occurrence proved to be troublesome as it suppresses abundance and richness of numerous native species (Chmura, 2013; Woziwoda *et al*., 2014a). For both oaks, the primary means of reproduction is thought to be abandonment of seed caches made by scatter-hoarding rodents and birds (den Ouden *et al*., 2005; Steele, 2008; Myczko *et al*., 2014). Nonetheless, whether caching benefits exceeds the costs imposed by seed predation has never been experimentally evaluated.

Drawing on past studies, we evaluated the following hypotheses (Table 1): (1) Benefits of seed caching should be larger than transportation benefits. Seed burial plays a critical role in regeneration of many plant species because it protects seeds from desiccation and strict seed predators (Haas & Heske, 2005; Vander Wall, 2010; Zwolak *et al*., 2016). On the other hand, rodents move seeds over rather short distances (within population: den Ouden et al. 2005), which is unlikely to result in strong distance-dependent effects. (2) The distance-related benefits should be smaller in red oak than in sessile oak. This result might occur because invasive species often escape from the inhibitory effects of soil biota in their alien range (Agrawal *et al*., 2005; Maron *et al*., 2014). (3) Seed removal by rodents should increase with distance from adult plants. This result might occur because dense seed shadows under the tree may create local satiation effects (Xiao *et al*., 2015). (4) Pilferage of cached acorns should be more frequent in red than in sessile oaks. This result might occur because heavier acorns are more likely to be pilfered (Perea *et al*., 2016) and red oak acorns are over twice heavier than sessile oak acorns (Bogdziewicz *et al*., 2017a). (5) Benefits of seed dispersal and burial will be large enough to compensate predation, leading to mutualistic relationship in which small mammals will aid recruitment of focal oaks. This prediction is based on results of a meta-analysis of previous studies, which suggested that outcomes of plant-rodent interactions tend to be weakly mutualistic (Zwolak and Crone 2012).

**Table 1.**
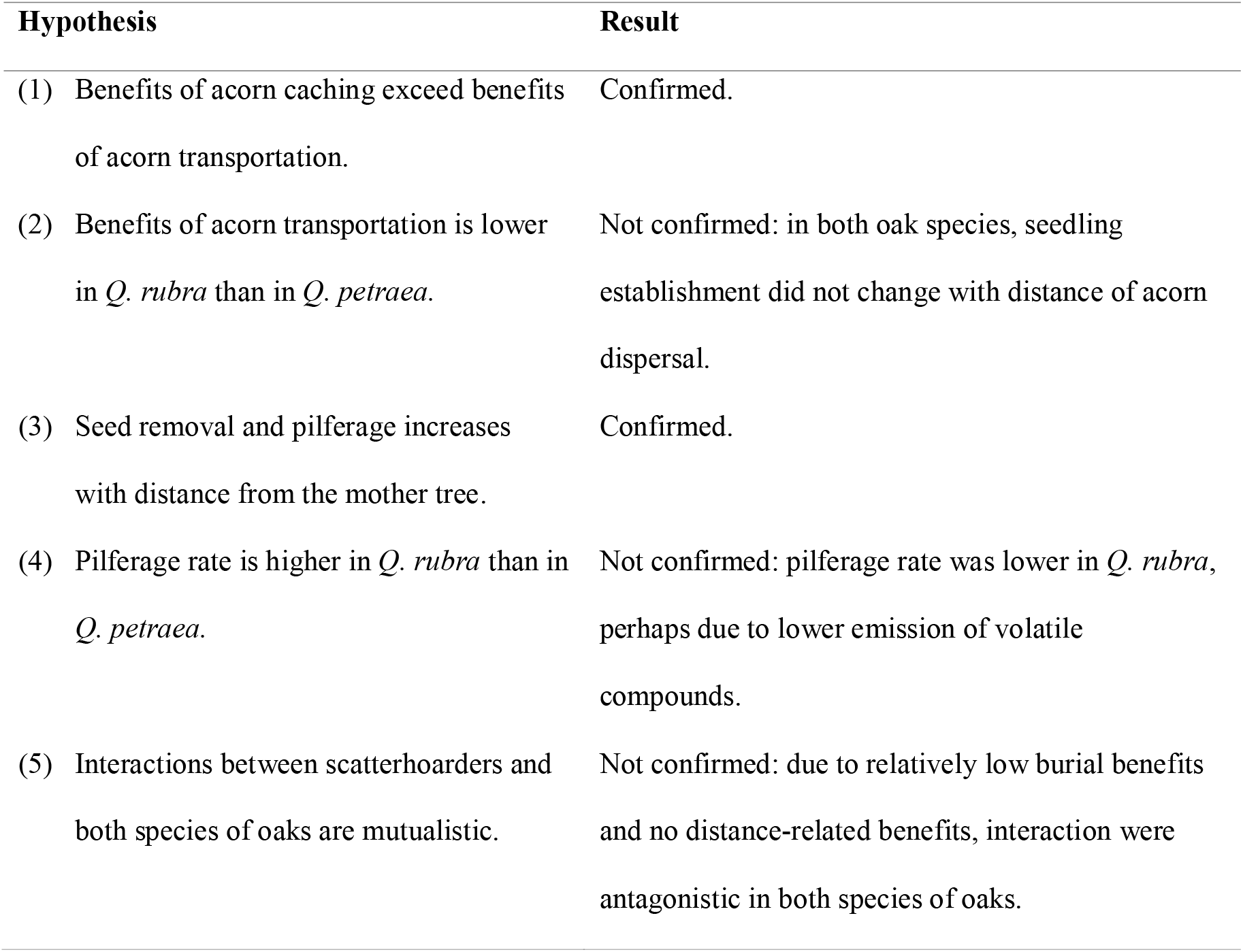
Summary of hypotheses and results.

We used the estimates of seedling establishment probability obtained in field experiments to place the focal interactions at the antagonism-mutualism continuum, using the modeling framework developed by Zwolak and Crone (2012). However, the original model did not include the potential hanges in caching benefits driven by seed pilferage (Steele *et al*., 2014; Sunyer *et al*., 2015; Zwolak *et al*., 2016). Therefore, as part of this paper, we extend the logic of the *p_c_* calculation to include pilferage by naïve foragers.

## Materials and Methods

We established four study sites in sessile – red oak mixed forests in Drawieńska Forest, western Poland (spaced 1 – 15 km from each other). This area is located in the temperate climate zone, with average annual precipitation of 592 mm and means monthly temperature ranging from 17°C in July to −2°C in January. These mixed forests comprise almost exclusively of the three oak species (Q. *petraea, Q. robur*, and *Q. rubra)*, with single trees of common hornbeam *(carpinus betulus)* and European beech *(Fagus sylvatica)*. The understory is poorly developed, with some patches of raspberry *(Rubus* sp.) and common nettle *(Urtica dioica)*, and seedlings of oaks and beech. Oak acorns are relatively large (average weight; *Q. petraea* 1.26 g., *Q. rubra:* 2.85 g.), and readily dispersed and eaten by small mammals (Bogdziewicz et al. 2017b, Bogdziewicz, unpublished). As revealed by camera traps, small mammals at our study sites were dominated by *Apodemus* sp., most likely *A. flavicollis*, a seed specialist (Selva *et al*., 2012, Gasperini *et al*., 2017).

To quantify the effects of acorn burial on seedling emergence we conducted seed addition experiments. We randomly chose 12 *Q. petraea* trees (3 per site) and 12 *Q. rubra* trees (3 per site). We added acorns of the focal species in 20 x 20 x 20 cm wire mesh cages (5 acorns per cage). Cages were buried 10 cm into the ground in sets of four. In half of the cages we buried acorns 1-2 cm into the ground and in the other half we placed acorns at the top of the litter layer and covered them with leaves to mimic autumn leaf fall. This treatment was crossed with rodent exclusion: in half of the cages we cut 8 x 8 cm holes to allow rodent access, and the other half remained closed to exclude rodent foraging. A comparison of seedling recruitment from acorns that were buried vs. placed on surface allowed us to estimate burial-dependent benefits of rodent seed dispersal. Rodent exclusion allowed us to estimate seed pilferage by comparing recruitment of buried acorns in open vs. closed cages.

To address the distance-related effects of rodent seed dispersal, the above-described cage sets were placed along transects. Under each tree, we established a transect along one cardinal direction, aiming to maximize the distance of the transect to other conspecifics. This was done assuming that rodents tend to carry and cache seeds towards areas of lower conspecific seed density (Stapanian & Smith, 1984; Hirsch *et al*., 2012; Steele *et al*., 2014; Yang *et al*., 2016). Thus, our estimates of distance-dependent effects may be overly positive, if such directed dispersal does not occur in our system. We placed five sets of cages at each transect. We used tree crown as a reference point and buried one set of cages directly underneath the crown border, another set 5 m towards the tree trunk (underneath the crown), and the remaining 3 sets every 5 m in the opposite direction (i.e. away from the tree trunk). We used 25 m as the maximum evaluated distance because acorn-tracking experiments report that vast majority of acorns transported by rodents are cached within that radius (den Ouden *et al*., 2005; Xiao *et al*., 2005; Muñoz & Bonal, 2011; Bogdziewicz *et al*., 2017b). We set up experimental cages in October 2016 and quantified seedling establishment in August 2017. The overall sample size equaled 2400 acorns (480 seedling cages).

### Statistical analysis

To test how acorn burial, distance from the tree, and rodent foraging affect the seedling establishment, we built a separate generalized linear mixed model (GLMM) for each oak species. We used nested random effects of cage set, tree, and study site, logit link, and binomial family error distribution, and implemented the models via lme4 package in R (Bates et al. 2015). In each model, we used proportion of established seedlings as the response variable, and burial (surface vs. sowed), rodent access (excluded vs. allowed), and distance from the tree as fixed effects. We also included all possible 2way interaction terms between fixed effects, and the 3-way interaction (which was removed when non-significant). We calculated marginal (i.e. the proportion of variance explained by fixed effects) and conditional (i.e. the proportion of variance explained by fixed and random effects) R^2^ for GLMMs using the MuMIn package (Nakagawa & Schielzeth, 2013; Bartoń, 2016).

Calculating the *p_c_* threshold and the effects of seed pilferage We evaluated how the interactions between rodents and oaks are placed along the antagonism–mutualism continuum (Zwolak & Crone, 2012). The *p_c_* threshold was calculated as a ratio of emergence from seeds sown on surface vs. emergence from buried seeds, with rodents excluded. Seed pilferage was gauged as the ratio of seedling recruitment from buried seeds in open vs. closed cages. Implicitly, the original definition of the proportion of seeds cached and uneaten (*p_c_*) combined three processes (Zwolak & Crone, 2012): the probability that a seed is cached, the probability it is eaten by the cache owner, and the probability it is pilfered:

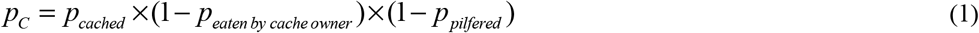

where *p_cached_* is the probability a seed is cached, *p_eaten_* by cache owner is the probability of retrieval by individuals responsible for seed burial and *p_pilfered_* is the probability of retrieval by pilferers. The *p_c_* threshold is the minimum value of when the benefits for plants balance the costs of seed consumption. Thus, if the threshold is determined by the proportion of seedling emergence from surface, *e_s_* (estimated with data on seedling emergence from seeds sown on surface in closed cages), to emergence from caches, *e_c_* (i.e. by benefits of burial, estimated with data on seedling emergence from seeds buried in closed cages), i.e.:

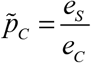

then the equation can be expanded to show the effects of pilferage:

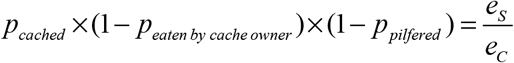

and rearranged to include only the unknown proportion of seeds cached and uneaten by the cache owner:

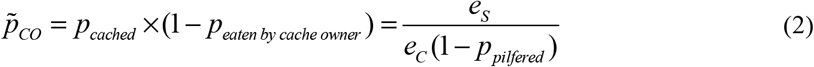

In the above equation, *p_CO_* is the minimum beneficial proportion of seeds cached and uneaten by *cache owner*, and all other parameters are as defined above. In other words, *p_co_* threshold defines the burial benefits while taking into account cache pilferage by naïve forages.

Confidence intervals for these parameters (*e_s_ e_c_*, and *p_c_*) were obtained with parametric bootstrapping, i.e. sampling from the distributions defined by the mean and standard error of each coefficient to obtain a joint distribution for the derived variables. We repeated the calculations of *p_c_* and *p_CO_* for both near the conspecific probability of germination (i.e. germination rate estimated at the distance 0 m, and far i.e. germination rate estimated at distance 25m). Probabilities of germination were derived from the above-described GLMMs.

The empirical *p_CO_* value (the ultimate probability that the acorn will be cached and not retrieved, accounting for retrieval by the whole granivore community) for both oak species was derived from a parallel study investigating rodent seed dispersal of the focal oaks, i.e. 17% for red oaks and 2% for sessile oaks (Bogdziewicz et al. unpublished).

## Results

In accordance with hypothesis (1), benefits of acorn burial were higher than benefits of acorn transportation away from adult trees. Acorn burial enhanced seedling establishment in both species, mainly through reducing seed predation. In *Q. petraea*, when rodents had access, acorn burial increased establishment probability 2-fold (open cages, surface vs. buried acorns: 18% vs. 39%). This effect was considerably weaker when rodents were excluded (closed cages, 57% for acorns on the surface vs. 65% for buried acorns; rodent exclusion × burial interaction in Table 2a, Fig. 1). Similarly, burial increased establishment probability almost 3-fold in *Q. rubra* (open cages: 25% vs. 71%, Fig. 1). This effect was weaker when rodents were excluded (50% vs. 73%; rodent exclusion × burial interaction in Table 2b).

**Figure 1.**
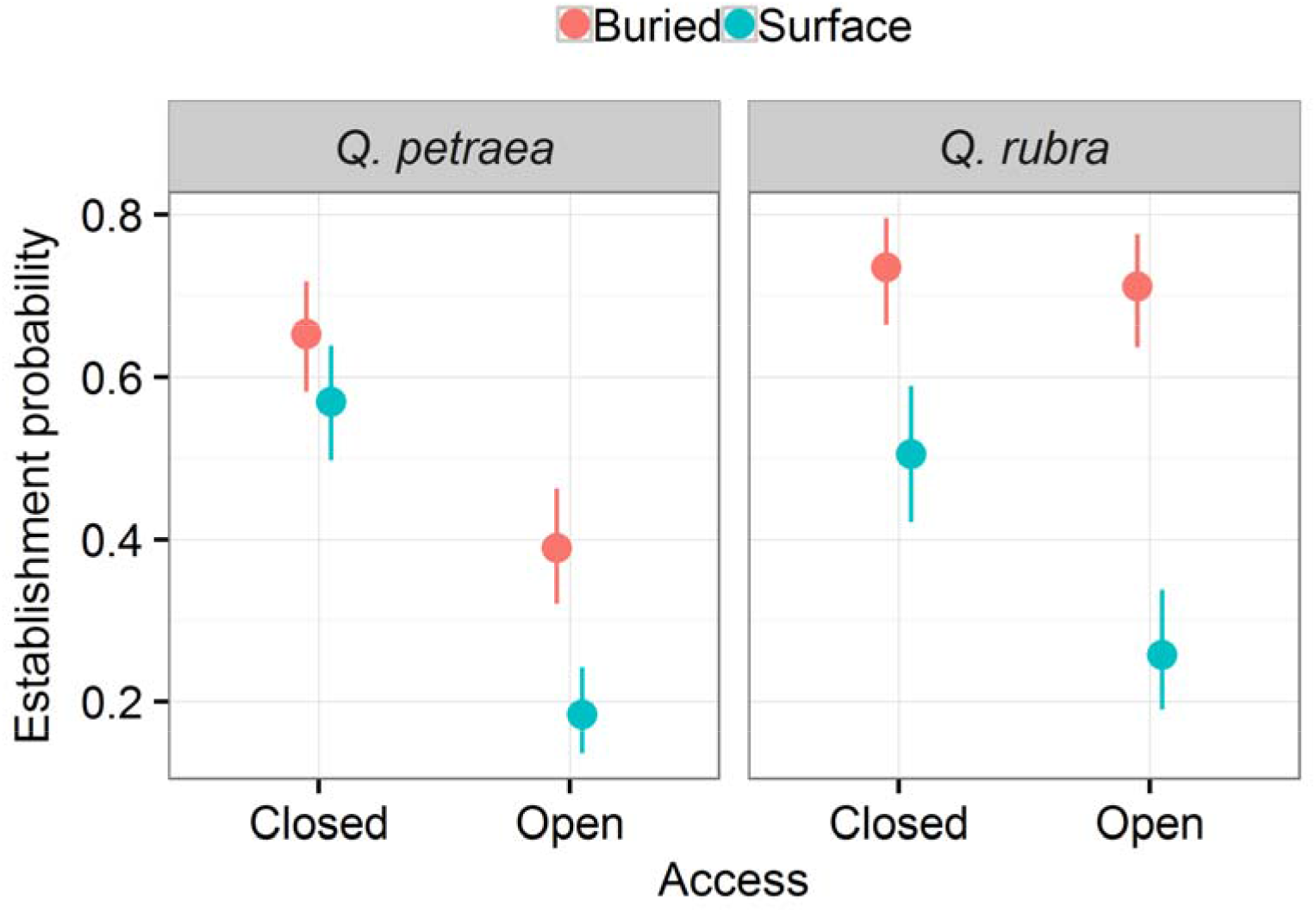
Probability of seedling establishment of *Q. petraea* and *Q. rubra* in closed (rodent access excluded), and open (rodent access allowed) cages. Sowed indicate acorns sowed in 1-2 cm in the soil, while surface indicate acorns sown on soil surface and covered with leaves to mimic autumn leaf fall. Whiskers indicate standard error.

**Table 2.**
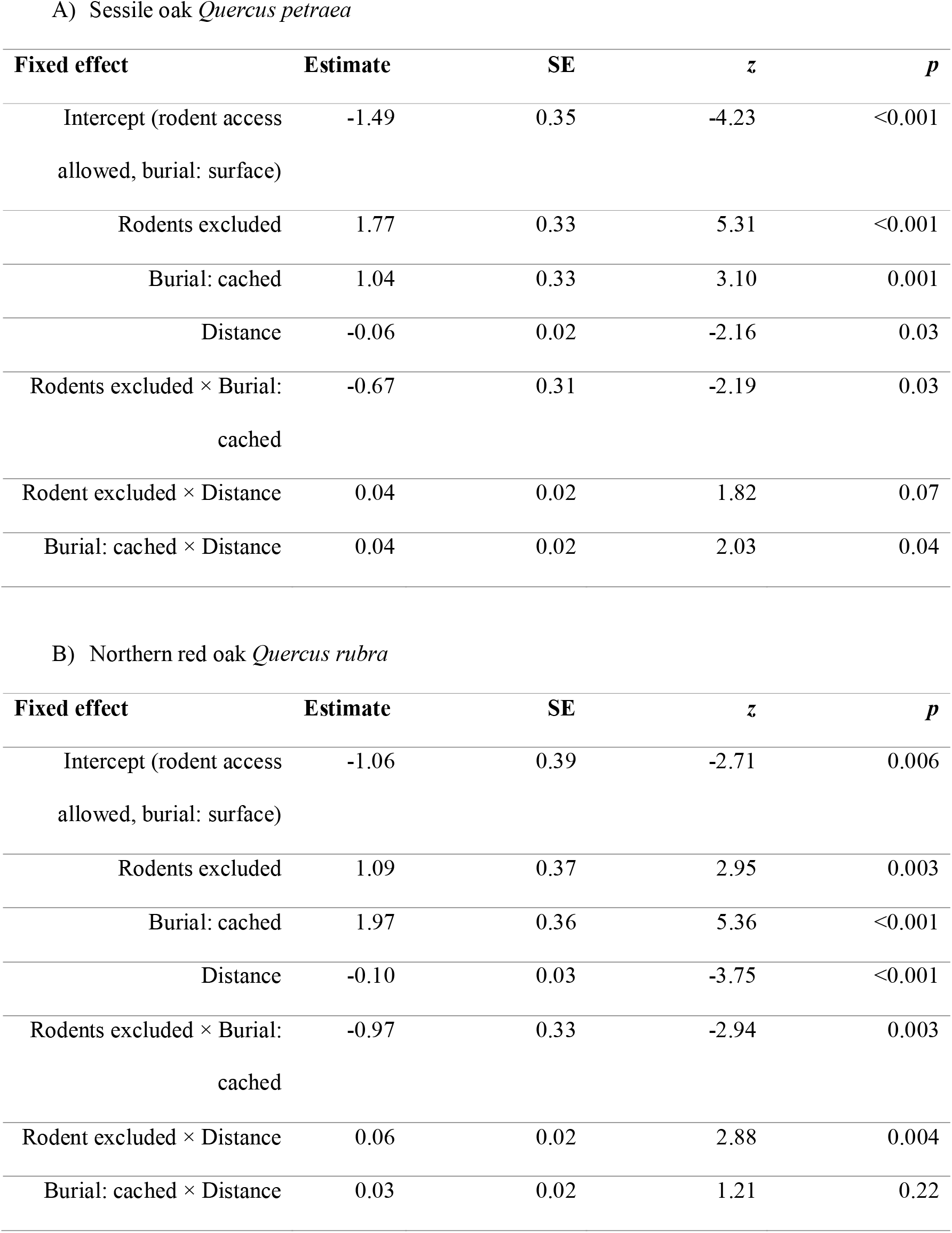
Effects of burial, distance from the parent tree, and rodent access on the germination probability of focal oaks. The marginal R^2^ of the model for *Q. petraea* equaled 0.28, while conditional 0.34. For *Q. rubra*, the marginal R^2^ of the model equaled 0.33, while conditional 0.38.

| Fixed effect | Estimate | SE | *z* | *p* |
| --- | --- | --- | --- | --- |
| Intercept (rodent access allowed, burial: surface) | -1.49 | 0.35 | -4.23 | <0.001 |
| Rodents excluded | 1.77 | 0.33 | 5.31 | <0.001 |
| Burial: cached | 1.04 | 0.33 | 3.10 | 0.001 |
| Distance | -0.06 | 0.02 | -2.16 | 0.03 |
| Rodents excluded × Burial: cached | -0.67 | 0.31 | -2.19 | 0.03 |
| Rodent excluded × Distance | 0.04 | 0.02 | 1.82 | 0.07 |
| Burial: cached × Distance | 0.04 | 0.02 | 2.03 | 0.04 |

B) Northern red oak *Quercus rubra*
| Fixed effect | Estimate | SE | *z* | *p* |
| --- | --- | --- | --- | --- |
| Intercept (rodent access allowed, burial: surface) | -1.06 | 0.39 | -2.71 | 0.006 |
| Rodents excluded | 1.09 | 0.37 | 2.95 | 0.003 |
| Burial: cached | 1.97 | 0.36 | 5.36 | <0.001 |
| Distance | -0.10 | 0.03 | -3.75 | <0.001 |
| Rodents excluded × Burial: cached | -0.97 | 0.33 | -2.94 | 0.003 |
| Rodent excluded × Distance | 0.06 | 0.02 | 2.88 | 0.004 |
| Burial: cached × Distance | 0.03 | 0.02 | 1.21 | 0.22 |

We did not detect any distance-related benefits of seed dispersal and contrary to hypothesis (2), the oak species did not differ in this regard. In fact, seedling establishment probability decreased with the distance from the focal tree (Table 2, Fig. 2 and 3). This phenomenon was caused by an increase in acorn removal, which supports hypothesis (3). In *Q. petraea* increased removal was apparent only when acorns were sown on surface (Table 2a, Fig. 2). Pilferage of cached acorns did not differ with distance from the tree in this species. In *Q. rubra*, this effect occurred both for acorns that were buried and those that were left on surface.

**Figure 2.**
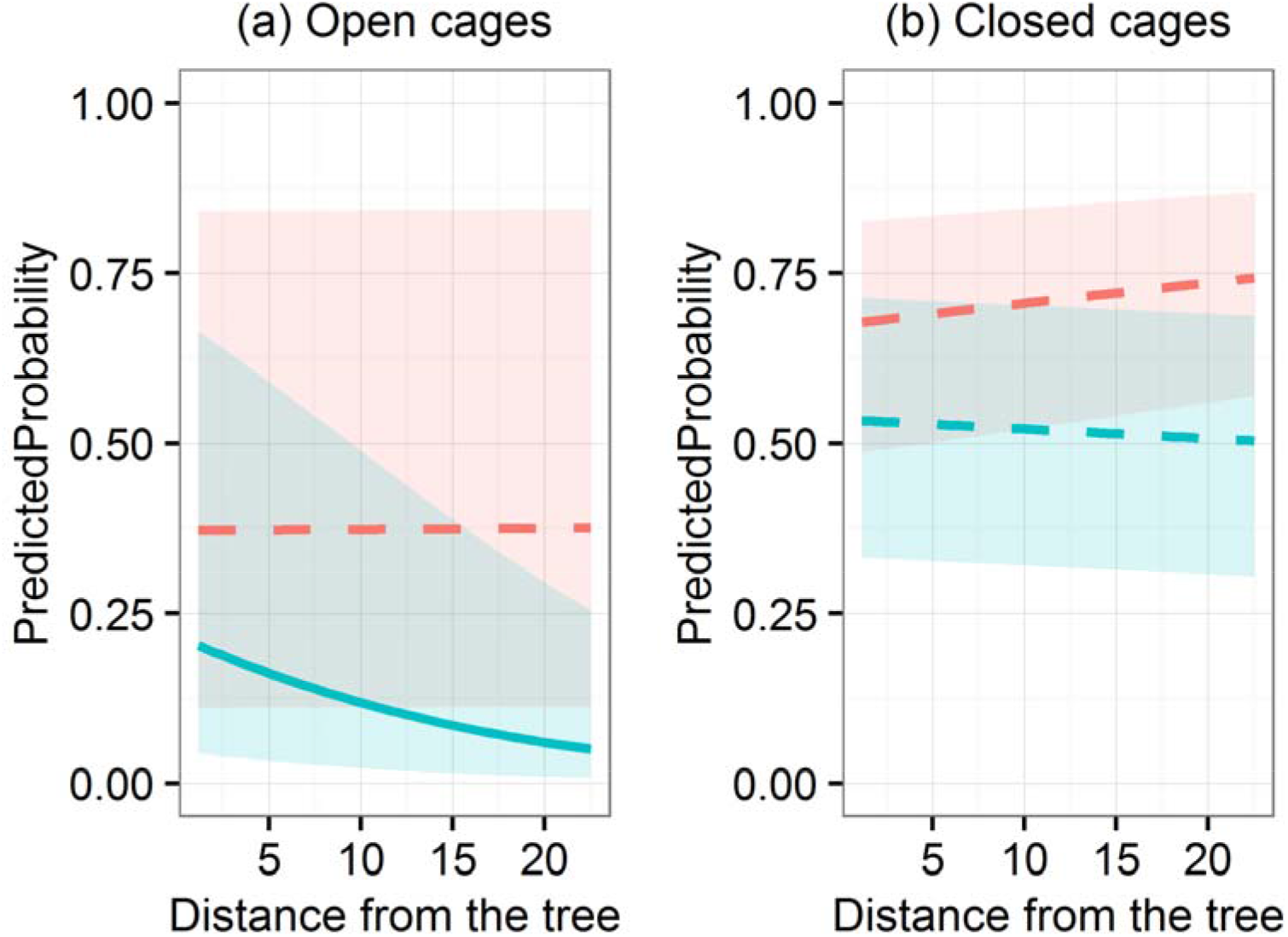
Probability of seedling establishment of sessile oaks *(Quercus petraea)* as a function of distance from the mother tree. Open cages indicate rodent access, while closed rodent exclusion; red lines acorns buried in topsoil, blue sowed on soil surface. Solid line indicates relationship significantly different from 0, while dashed line non-significant relationship. Shaded areas indicate 95% confidence intervals.

**Figure 3.**
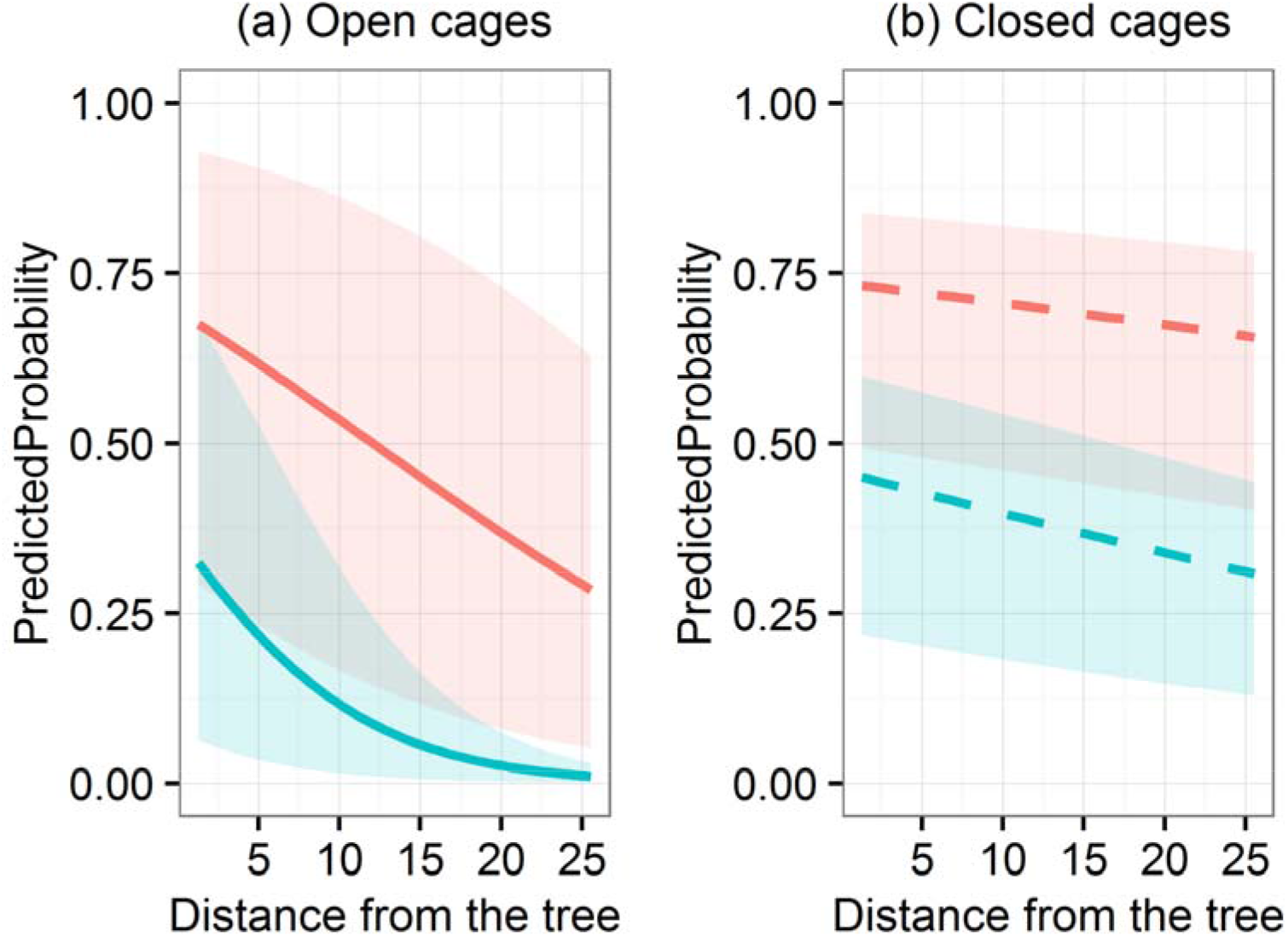
Probability of seedling establishment of red oaks *(Quercus rubra)* as a function of distance from the mother tree. Open cages indicate rodent access, while closed rodent exclusion; red lines acorns buried in topsoil, blue sowed on soil surface. Solid line indicates relationship significantly different from 0, while dashed line non-significant relationship. Shaded areas indicate 95% confidence intervals.

Acorn burial was more beneficial for *Q. rubra* than for *Q. petraea* (see above, and Fig. 1). In contrast to hypothesis (4), pilferage rates were higher in *Q. petraea* than in *Q. rubra*. In *Q. petraea* seedling establishment from buried acorns was 1.5 times higher when rodents were excluded (39% in open vs. 65% in closed cages). In *Q. rubra*, burial provided almost complete protection from pilferage (open vs. closed cages: 71% vs. 73%). Note that the percentages estimates are the model intercepts, and decrease with distance from the tree in some treatments (see below).

Estimated *p_CO_* values (the minimum beneficial proportion of seeds cached and uneaten *by cache owner*, i.e. those taking into account cache pilferage) equaled 1.21 (95% CI: 0.86-1.79) for *Q. petraea* (both near and far from the tree), and 0.69 (95% CI: 0.46-0.97) in *Q. rubra* near, and 1.24 (95% CI: 0.77-1.99) in *Q. rubra* far from the tree (Fig. 4). Note that the *p_CO_* value for *Q. petraea* does not differ with distance because the pilferage rates were distance-independent (Fig. 2a). These values are either impossible to reach (when they exceed 1) or would require almost all cached acorns to be never retrieved to approach the mutualism parameter space of the interaction (Fig. 4). Thus, hypothesis (5) was not supported: interactions between scatterhoarders and oaks in our study system were clearly antagonistic.

**Figure 4.**
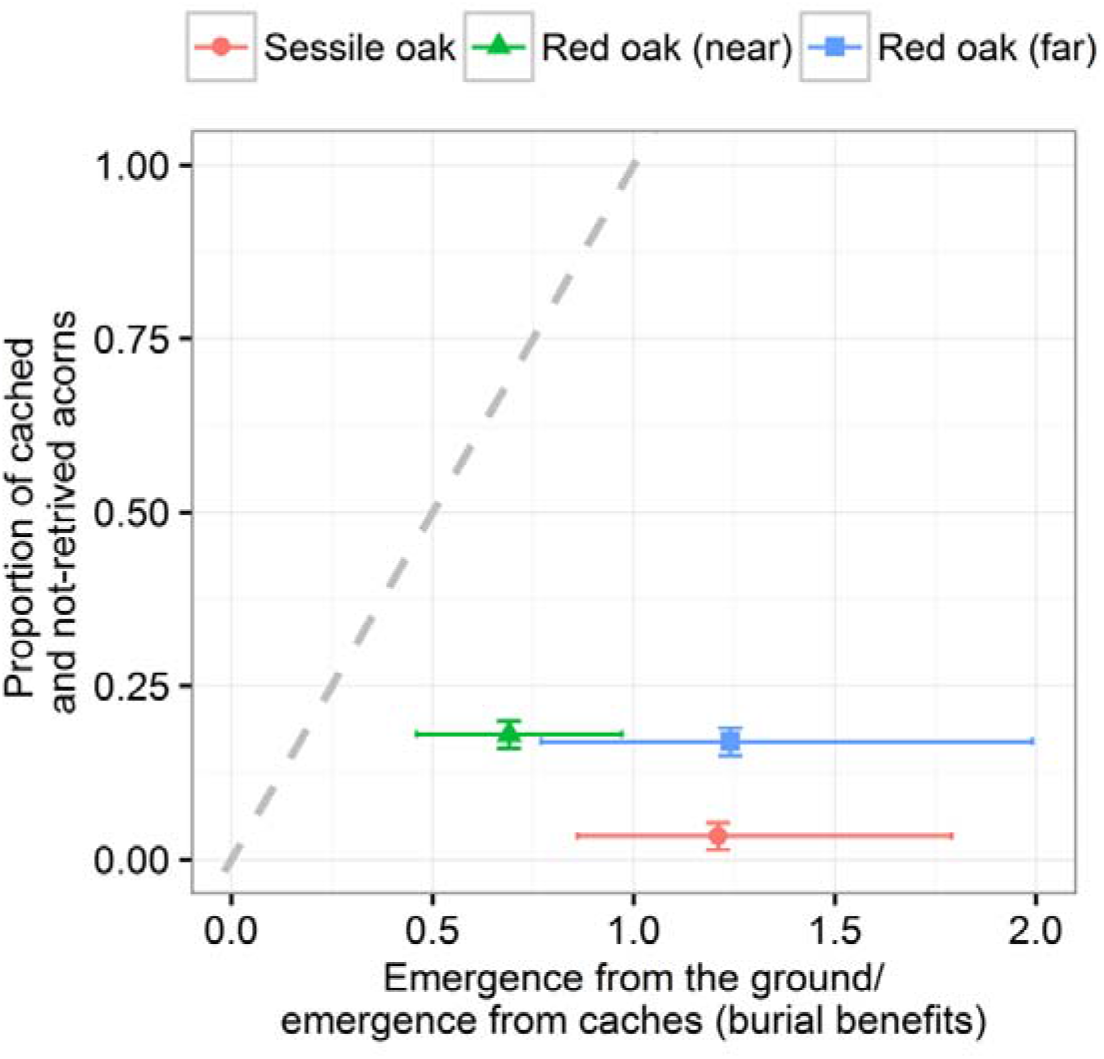
Classification of oak–granivore interactions based on the probability of caching and not retrieving seeds, and the ratio of seedling emergence from the ground to emergence from caches. The net effect of granivores is beneficial at any point above the dotted gray line and antagonistic at any point below. The ‘far’ and ‘near’ categories indicate the establishment ratio calculated based on germination rate estimated at the distance 0 m (near), and 25 m (far). For sessile oaks, the ratio components did not differ with the distance from the seed source tree (see Fig. 2). The values on x-axis (proportion of acorns cached and not retrieved) are derived from parallel study (Bogdziewicz et al. unpublished).

A fraction of removed acorns could be recached rather than consumed. If we assume that pilfered seeds are as likely to be eaten or re-cached as seeds collected for the first time (Jansen *et al*., 2004), then the consequences of burial depend on the number of rounds of recaching (Online Supplement 1). Nonetheless, for the parameters observed in our system, recaching of pilfered acorns would not affect the conclusion that scatterhoarders acted antagonistically in their interactions with oaks (Online Supplement 1).

## Discussion

Our study is the first to separate and directly quantify two most important services provided to plants by their rodent partners: seed transportation away from parent plants and seed burial in topsoil. Our results suggest that widely-accepted benefits of transportation might in many ecological systems be smaller than expected. Moreover, they demonstrate that even relatively large improvements in seedling establishment after seed burial do not necessarily render the plant-scatterhoarder interaction mutualistic. Finally, and most generally, our study illustrates a straightforward empirical approach that can be used to evaluate the ambiguous role of scatterhoarding granivores in plant regeneration.

This approach consists of quantifying the net outcome of an interaction through assembling different parameters piecewise with a simple mathematical model. This method, although used here to a specific plant-scatterhoarder dataset, is very general: in fact, it is analogous to building population models with separately-measured vital rates (e.g. Caswell 2001, Morris and Doak 2002). Similarly, just like other ecological models, it is a simplification, that can be made more realistic by adding additional information (see, e.g., Maron and Crone 2006, Ehrlen et al. 2016 for examples in a demographic context). Here we used data on probability of seedling recruitment with and without rodents; near and far from adult conspecifics; with and without seed pilferage. Possible future extensions of our seed-scatterhoarder model include e.g. effects of temporal variability or effects of directed dispersal of seeds into particularly favorable microsites (see below).

In addition, this case study provides a method to compare the importance of different types of benefits or costs of interspecific interactions. Animal dispersers often provide several different services that are not of equal value for their partner plants, but benefits provided by each are rarely separated. Yet, a few case studies that have done so demonstrated that it may change the way we think about particular interactions. For example, birds that disperse chili peppers *(Capsicum chacoense)* remove pathogens from the dispersed seeds (condition-related benefit) and transport seeds far from parent plants (escape-related benefit). However, only gut passage enhances seed survival (Fricke *et al*., 2013). A similar situation was reported for Iberian pears *(Pyrus bourgaeana)*, where pulp removal was more important than transportation distance for plant recruitment (Fedriani *et al*., 2012). By untwining the role of caching (condition-related benefit) from transportation distance in oak-scatterhoarders interactions, we demonstrated that acorn burial was the main benefit of this interaction (hypothesis 1 in Table 1), at least during the period of the study. The lack of distance-dependent benefits was unsurprising because in locally common plant species, the benefits should disappear when the species become so common that their predator and pathogen communities become functionally uniform across the landscape (Janzen, 1971; Schupp, 1992; Fricke *et al*., 2013; Garzon-Lopez *et al*., 2015). This is likely the case in our system because sessile and red oaks dominate forest stands. Our results suggest that the generally greater establishment of some species far from parent plants may be due distance-independent benefits of burial rather than distance-dependence *per se*. In other words, intertwined caching and transportation may create false positive effect of transportation, while it is only caching that helps recruitment (as in our system). This calls for increased attention on condition-dependent benefits of seed dispersal, which have been often overlooked as researchers focused on dispersal distance and final location of seeds (Fricke *et al*., 2013).

The patterns of seedling establishment suggest that rodent foraging is a strong filter of oaks spatial recruitment in our system. When rodents were excluded, distance from the tree did not change the probability of seedling establishment in either species (contrary to hypothesis 2 in Table 1). However, when rodent foraging was allowed, seedling establishment rate was highest near the trees, indicating that locally dense seed shadows may allow higher proportion of undispersed seeds to survive and germinate (hypothesis 3 in Table 1). On the other hand, distance to adult trees influenced pilferage of buried acorns in a species-specific manner. In *Q. petraea*, the pilferage did not change with distance from the tree, while it decreased with distance in *Q. rubra*. It suggests that ambient seed density has a stronger effect on cache pilferage rates in *Q. rubra* than in *Q. petraea* (Gálvez *et al*., 2009). However, we did not measure seed shadows, and thus this pattern might have resulted from larger crop size of *Q. rubra* at the time of our experiments.

*Q. rubra* acorns were generally better protected from seed predation once buried than acorns of *Q. petraea* (contrary to hypothesis 4 in Table 1). The pilferage rates of buried seeds are related to their release of odorant molecules (Vander Wall, 1998; Vander Wall & Joyner, 1998; Yi *et al*., 2016). *Q. rubra* appears to rely more on rodent dispersal in their native range than does *Q. petraea* in Europe (den Ouden *et al*., 2005; Steele, 2008). Thus, the tighter coevolution between small mammals and *Q. rubra* in the species native range (Steele, 2008), may led to reduced emission of volatile compounds in this species, reducing the overall pilferage rates. In consequence, lower pilferage rates in *Q. rubra* increased overall burial benefits, moving the interaction closer to the mutualism.

Nonetheless, in contrast to our fifth and final hypothesis (Table 1), scatterhoarding rodents reduced recruitment of focal oak species during the period of our study. As anticipated, acorn burial increased the likelihood of seedling establishment. However, the seedling establishment of unburied acorns in the absence of small mammal foraging was high for both species. As a final result, burial benefits were too small to override the costs of seed predation (Zwolak & Crone, 2012). In fact, even if the probability of seedling establishment from rodent caches would elevate to 100%, the *p_CO_* value for *Q. petraea* would equal 0.57 while for *Q. rubra* 0.50. This level of survival of cached acorns appears unlikely, as the reported values range from 1 to 20% (Vander Wall & Joyner, 1998; Vander Wall, 2002; Gomez *et al*., 2008; Xiao *et al*., 2013; Bogdziewicz *et al*., 2017b; Wróbel & Zwolak, 2017). Therefore, while it is still best for an acorn to be cached and then forgotten, presence of small mammals did not help recruitment of oaks in our system.

We note, however, that the balance of benefits and costs in conditional mutualisms typically changes over time (Theimer 2005, Klinger and Rejmánek 2010, Zwolak et al. 2016) and our study was conducted during relatively short time frame (1 year). It is possible that fluctuating environmental conditions (e.g. years with droughts or severe winters) can increase the benefits of acorn caching and shift the oak-rodent relationship towards mutualism. This is particularly likely given that burial benefits in our study were remarkably low: e.g. Kollmann & Schill (1996), García *et al*. (2002), and Xia *et al*., (2016) reported higher benefits in oaks. In this situation, the role of small mammals in oak recruitment would change over time. Moreover, both focal oaks species exhibit mast years (Sork *et al*., 1993; Bogdziewicz *et al*., 2017c), which in turn drives fluctuations in small mammal population abundance and may create satiation effects, both at the seed source and after caching (Kelly, 1994; Xiao *et al*., 2013; Zwolak *et al*., 2016). Yet, the potential increase in cache survival caused by the phenomenon is unlikely to counterbalance the costs of seed predation – unless it is accompanied by environmental changes that increase the benefits of seed caching. Thus, an interesting venue for future studies would be to quantify temporal variation in this interaction.

As a final caveat to this study, Janzen-Connell effects are stronger at the seedling than seed-to-seedling stage (Comita *et al*., 2014). Therefore, the benefits of transportation may appear at later stages of plants life cycle. However, several studies that evaluated distance-dependent survival rates at seedling stage in temperate oaks did not found the effect (Reinhart *et al*., 2012; Comita *et al*., 2014). Furthermore, directed dispersal increases the likelihood of colonization of microhabitats that are favorable for germination and establishment (Steele *et al*., 2013; Yi *et al*., 2013). Although such effects were never reported for focal oaks, our experimental design could underestimate these effects and thus, benefits of acorn transportation by scatterhoarding rodents.

To conclude, we presented simple means by which outcomes of conditional plant-scatterhoarder interactions can be classified. The strength of our approach lies in its versatility: it uses mathematics to combine different types of data and can be easily modified to incorporate new information when data on other parameters becomes accessible. Our empirical results demonstrated that certain common assumptions (e.g. that scatterhoarding by rodents invariably improves plant recruitment; that improved germination after seed burial is sufficient to make plant-scatterhoarder interactions mutualistic; that transportation away from maternal plants is highly beneficial) do not always hold and should be tested rather than taken for granted.

## Acknowledgements

We thank Nadleśnictwo Krzyż Wlkp. for the permission to conduct the study. Kinga Stępniak, Agnieszka Amborska-Bogdziewicz, and Jan Bogdziewicz helped during the field work, and Anna and Bogusław Bogdziewicz provided logistical assistance. Dawid Manyś provided invaluable assistance during field site selection. The study was supported by the (Polish) National Science Centre grant ‘Preludium’ no. 2015/17/N/NZ8/01565.

